# Profile likelihood-based parameter and predictive interval analysis guides model choice for ecological population dynamics

**DOI:** 10.1101/2022.09.05.506701

**Authors:** Matthew J Simpson, Shannon A Walker, Emma N Studerus, Scott W McCue, Ryan J Murphy, Oliver J Maclaren

## Abstract

Calibrating mathematical models to describe ecological data provides important insight via parameter estimation that is not possible from analysing data alone. When we undertake a mathematical modelling study of ecological or biological data, we must deal with the trade-off between data availability and model complexity. Dealing with the nexus between data availability and model complexity is an ongoing challenge in mathematical modelling, particularly in mathematical biology and mathematical ecology where data collection is often not standardised, and more broad questions about model selection remain relatively open. Therefore, choosing an appropriate model almost always requires case-by-case consideration. In this work we present a straightforward approach to quantitatively explore this trade-off using a case study exploring mathematical models of coral reef regrowth after some ecological disturbance, such as damage caused by a tropical cyclone. In particular, we compare a simple single species ordinary differential equation (ODE) model approach with a more complicated two-species coupled ODE model. Univariate profile likelihood analysis suggests that the both models are practically identifiable. To provide additional insight we construct and compare approximate prediction intervals using a new parameter-wise prediction approximation, confirming both the simple and complex models perform similarly with regard to making predictions. Our approximate parameter-wise prediction interval analysis provides explicit information about how each parameter affects the predictions of each model. Comparing our approximate prediction intervals with a more rigorous and computationally expensive evaluation of the full likelihood shows that the new approximations are reasonable in this case. All algorithms and software to support this work are freely available as jupyter notebooks on [GitHub](https://github.com/ProfMJSimpson/profile_predictions) so that they can be adapted to deal with any other ODE-based models.

## 1. Introduction

Appropriate model choice is a difficult problem with, in our view, no universal or automatic solution. However, *identifiability analysis* [6, 14, 22, 25, 33–35, 46] provides one set of tools that can help guide model choice in terms of determining an appropriate level of model complexity relative to available data. Formally, a model is *identifiable* when distinct parameter values imply distinct distributions of observations [25]. Less formally, a model is identifiable when it is possible to uniquely determine the model parameters using an infinite amount of ideal data. While a model being non-identifiable does not mean the model is wrong, it is, as a whole, then essentially unfalsifiable, although some aspects of the model may still be falsifiable or otherwise testable. Thus we may decide to prioritise the use of identifiable models over non-identifiable models. Often in the systems biology literature, *identifiability* is also referred to as *structural identifiability* [33–35, 46], and working within the framework of ordinary differential equation (ODE)-based models, it is possible to formally analyse the structural identifiability using a range of algebraic methods [3, 8, 9, 24]. This kind of analysis can be very insightful and aid in the choice of identifiable models over non-identifiable models, or help reformulate non-identifiable models into simpler identifiable models [12, 26, 27].

The focus of the present work is to interpret a set of real ecological field data using different mathematical models framed in terms of ODEs. Specifically, the data we consider describes the temporal increase in the percentage coral cover on a coral reef after some kind of natural disturbance, such as a tropical cyclone [49]. Such coral recovery data can be interpreted in several different ways, and there is no standard approach to interpreting this data. The simplest approach is to work with the evolution of the total hard coral cover as a function of time. Another is to consider dividing the total hard corals into different coral groups and consider modelling the temporal evolution of each group of hard corals. In essence, these approaches amount to working with either: (i) a single ODE model for the total hard coral cover or; (ii) a system of coupled ODE models describing the coral cover associated with each grouping. While structural identifiability analysis can aid model choice in general, here both sets of differential equations are already structurally identifiable [42] and the data are sparse and noisy. Thus we require additional means of guiding model choice and to assess the associated trade-off between model complexity and data availability. Here we focus on *practical identifiability*. While the formal definition of structural identifiability requires an infinite amount of ideal data, the less formal concept of practical identifiability (or *estimability*) seeks to describe whether it is possible to provide reasonably precise estimates using finite, non-ideal data [10, 25, 50]. This question typifies the kind of scenario one can expect to confront when using mathematical models to interpret biological and ecological data more generally.

In the literature, practical identifiability can be assessed in several different ways. In both Bayesian and (standard parametric) frequentist frameworks, identifiability is usually considered a property of the likelihood function, although a related but distinct Bayesian concept of *posterior identifiability* has also been proposed [10]. Computationally, practical non-identifiability can often be detected in a Bayesian setting through Monte Carlo Markov Chain (MCMC) sampling techniques when the samples fail to converge [17, 38, 39]. An alternative approach is to focus directly on the likelihood function and use profile likelihood and computational optimisation to construct profiles for each parameter, or for combinations of parameters [4, 14, 17, 31, 39]. Such profiles are routinely used to construct frequentist asymptotic confidence intervals for target parameters [31, 36]. Our previous work indicates that profile likelihood analysis can lead to quantitatively similar conclusions as, for example, MCMC-based analysis, with the main practical difference being the profile likelihood analysis is often far less computationally demanding [39]. Other authors have also emphasised the suitability of profile likelihood for practical identifiability analysis relative to other approaches such as MCMC, bootstrap, or Fisher information-based methods [14, 34]. Therefore, as part of our study, we carry out a profile likelihood-based analysis. The computational differences between using MCMC-based sampling methods compared to optimization-based profiling methods is most significant for more computationally demanding models, such as working with systems of nonlinear partial differential equations [39].

Working in a profile likelihood-based framework, we can assess models in terms of the widths of their associated parameter confidence intervals as an informal strategy for trading off model complexity with data availability. This provides a potentially more insightful means of model comparison relative to overly simplistic information criteria-based methods [1, 5, 20, 37, 42, 48]. However, in addition to constructing and comparing confidence intervals for each parameter in all models considered, we also construct parameter-wise *predictive* intervals for the mean coral cover trajectory using the univariate parameter profile likelihoods. (Note: here, we use the term *predictive* for any output from running the *forward* model given a set of parameters, that is we consider the mean trajectory as a predictive quantity.) This parameter-wise approach to prediction enables us to separate the roles of each parameter in terms of a more tangible model prediction rather than a less tangible parameter estimate [29]. For our models of interest, we show that generating parameter-wise predictions based on profile likelihoods provides an extremely efficient means of carrying out a sensitivity study and constructing approximate predictive intervals for the mean trajectory in the models we consider. Our approximate prediction intervals are analogous to performing a posterior predictive check in a Bayesian framework, except that our approach is likely to be more computationally efficient. This information provides insight that is complementary to the usual parameter confidence intervals considered in a typical practical identifiability analysis and helps guide model choice according to user goals, for example parameter inference or prediction or some combination of these.

Overall, this work provides insight into the question of model choice for scenarios where models are structurally identifiable, but may differ in terms of practical identifiability. Although we motivate this work with an ecological field data set describing coral reef regrowth after a disturbance, our approach can be applied directly to any model based on ordinary differential equations. All of our software is freely available in the open source Julia language on [GitHub](https://github.com/ProfMJSimpson/profile_predictions).

## 2. Field data and modelling objectives

We work with field data describing regrowth of hard corals off an island on the Great Barrier Reef, Australia. The regrowth process involves coral recolonisation after some external disturbance, such as the impact of a tropical cyclone [13]. Developing the ability to understand the mechanisms that drive this regrowth, and to predict the regrowth process, is extremely important as climate change-related disturbances increase in frequency [18, 49]. For example, if the time scale of regrowth is shorter than the time between disturbances we might anticipate that regrowth and recovery is likely, however if the time scale of regrowth is longer than the time between disturbances we might anticipate that regrowth and recovery is unlikely. One way to develop this understanding of the predictive capability is to calibrate a mathematical model to match the observations.

In this work we consider data describing the temporal regrowth of hard coral cover on a reef near Lady Musgrave Island, Australia (Figure 1(a)). Data in Figure 1(b) shows the temporal evolution of the total percentage area cover of hard corals, *S* (*t*) (blue discs). The field methods by which this data are collected are described by Warne et al. [48] and the data is available through the eATLAS repository [13]. In brief, data collection involved repeated human surveys of several sites near Lady Musgrave Island over a period of approximately 10 years after some external disturbance, such as a cyclone. The data presented in Figure 1 here comes from site 1, whereas the data presented in the Supplementary Material document comes from site 3. Data collection involves measuring the proportion of area covered by different types of coral species, and the reported data is averaged over many transects across each site. In this case, the data shows a classical sigmoid growth response with apparently exponential growth at early times, followed by a reduction in net growth as the total hard coral cover approaches some maximum density at later times [42, 49]. Data in Figure 1(c) shows more detailed information where we plot the time evolution of the percentage of the dominant hard coral group called *Acroporidae, C*_1_(*t*) (red diamonds), together with the time evolution of the percentage of all other hard corals grouped together, *C*_2_(*t*) (green hexagons). For these sites at Lady Musgrave Island the dominant hard coral group is *Acroporidae*, but this may not be the case on other coral reefs in different locations. We also plot the total hard coral cover, *S* (*t*) = *C*_1_(*t*) + *C*_2_(*t*) (blue discs), and note that the total hard coral cover in Figure 1(b) is identical to the total hard coral cover in Figure 1(c). Plotting the data in these two ways motivates the question of whether we should aim to model the time evolution of *S* (*t*) alone, or whether we should aim to model the time evolution of both *C*_1_(*t*) and *C*_2_(*t*). This is the main question we will explore through our model-based analysis.

**Figure 1:**
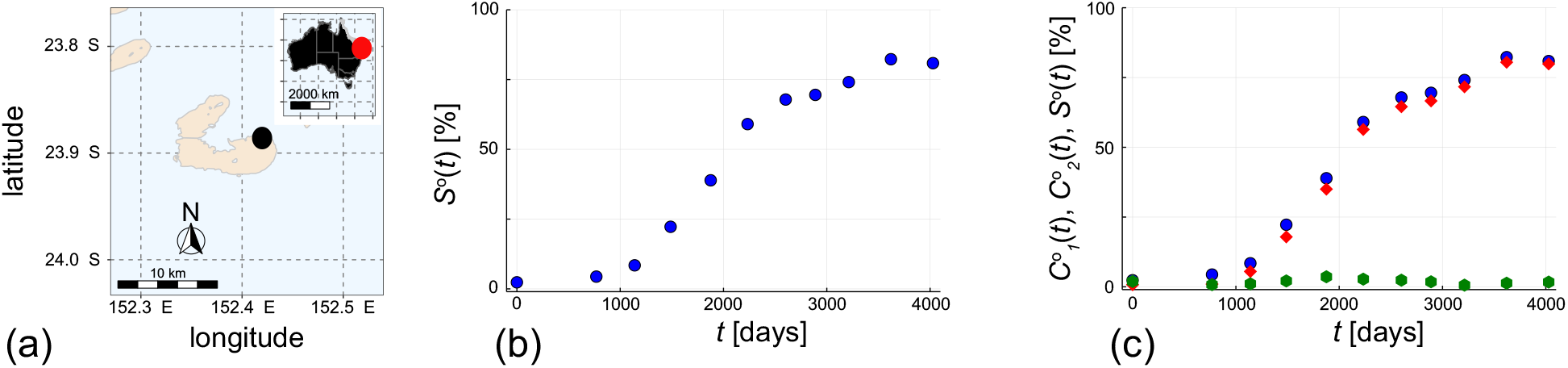
(a) Location of Lady Musgrave Island (black disc) relative to the Australian mainland (inset, red disc). (b) Field data showing the time evolution of the percentage total hard coral cover, *S* (*t*) (blue discs) after some disturbance at monitoring site 1. (c) Field data showing the time evolution of the percentage *Acroporidae* coral cover, *C*_1_(*t*) (red diamonds), the percentage cover of all other hard corals, *C*_2_(*t*) (green hexagon), and the total hard coral cover, *S* (*t*) = *C*_1_(*t*) + *C*_2_(*t*) (blue discs) after some disturbance at monitoring site 1. Note that the blue discs in (b) and (c) are identical. Time is measured in days after disturbance.

The field data in Figure 1 is associated with monitoring site 1 at Lady Musgrave Island. All algorithms and results used in this study are available on GitHub where we also present a second set of data for monitoring site 3. For completeness, all data and analysis associated with site 3 are given in the Supplementary Material documents. This additional data and the computational results for site 3 are presented in precisely the same format as given here for site 1. As we show, both data sets can be analysed with the same tools developed in this study, so we choose to present data for the site 1 here in the main document whereas the equivalent data and results for site 3 are provided on GitHub in the form of several user-friendly jupyter notebooks. It is worth noting the timescale of the coral regrowth in Figure 1 is approximately a decade, which is very different to a typical experiment in cell biology, for example, where cell proliferation takes places over a few days only [15, 19, 48]. Given the relatively rapid timescale of cell proliferation, it is typical for a cell biology experiment to be repeated several times to provide an estimate of the variability in the growth process [19, 48]. Here, in contrast, we deal with a very different type of biological growth process that takes approximately a decade, and so it is infeasible to repeat this experiment to provide a pre-estimate of the variability in the regrowth processes.

Throughout this work, all estimates of hard coral cover are expressed in terms of percentage coverage meaning that all model parameters associated with density are dimensionless. Further, the time-scale of regrowth is measured in days (Figure 1), so all growth and competition rates have dimensions of days^−1^.

## 3. Methods

### 3.0.1. Process models

As outlined in Section 1, we will model the data in Figure 1 using two different approaches. First we will use a very simple logistic growth model to describe the evolution of the total coral cover, *S* (*t*),

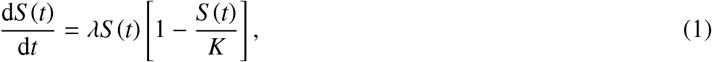

where *S* (*t*) is the percentage total coral cover at time *t, λ* > 0 is the growth rate and *K* > 0 is the long-time carrying capacity density. While Equation (1) can be solved exactly, we keep our approach general by solving all ODE models numerically, using standard numerical integration routines available in Julia [32]. The solution of Equation (1) depends upon the initial cover, *S* (0), which means when we solve (1) we must specify three model parameters: *λ, K* and *S* (0). While there are many other forms of single-species logistic growth-type models that we could use to capture this kind of sigmoid growth data [28, 45], our previous analysis of this data indicates that the logistic growth model performs better than alternative models (e.g. Gompertz model, Richards’ model) that encounter issues of practical identifiability [42].

The second approach we take is to divide the total population into two interacting subpopulations, *C*_1_(*t*) and *C*_2_(*t*), such that *S* (*t*) = *C*_1_(*t*) + *C*_2_(*t*), where *C*_1_(*t*) represents the percentage cover of the dominant hard coral *Acroporidae*, and *C*_2_(*t*) is the percentage cover of all other hard corals. We propose the following minimal, but realistic, model

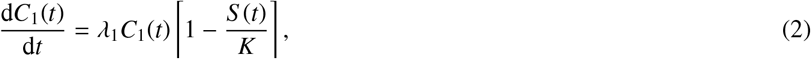

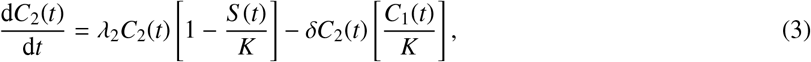

where *λ*_1_ > 0 is the low-density growth rate of *Acroporidae, λ*_2_ > 0 is the low-density growth rate of other hard corals, *K* > 0 is the long-time carrying capacity density and *δ* > 0 describes the rate at which *Acroporidae* competes with the other hard corals. This coupled model is constructed to capture the key features in the measured data since we see that the percentage of *Acroporidae* increases monotonically with time (Figure 1(c)), where as the percentage of other hard corals initially increases and then eventually decreases at later time (Figure 1(c)). These observations motivate the use of a simple logistic-like term governing the evolution of *C*_1_(*t*) in Equation (2), whereas the initial increase in other hard corals followed by the later decline in *C*_2_(*t*) motivates a combination of a logistic-like growth term and a competition term in Equation (3) as one straightforward way of capturing these trends. Of course, there are other potential terms that one could include in our model, and we will discuss some other options later in Section 5. However, we choose to work with this relatively straightforward form since it appears to capture the key features of the data and is ecologically plausible. Solutions of Equations (2)–(3) are obtained numerically [32] and require the specification of the initial coverage of both coral types, *C*_1_(0) and *C*_2_(0). Therefore, when we solve (2)–(3) we must specify six parameters: *λ*_1_, *λ*_2_, *K, δ, C*_1_(0) and *C*_2_(0).

### 3.1. Observation model

We assume that observed data 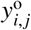 are measured at *I* discrete times, *t*_*i*_, for *i* = 1, 2, 3, …, *I*, and that at each time point we record *J* measurements, for *j* = 1, 2, 3, …, *J*. Within this framework, when we work with total hard corals we only have *J* = 1, whereas when we divide the total hard corals into *Acroporidae* and all other hard corals we have *J* = 2. We use a superscript ‘o’ to distinguish the noisy observed data from the model predictions. Regardless of whether we work with Equation (1) or (2)–(3), noise-free model predictions are denoted *y*_*i, j*_(*θ*) = *y*(*t*_*i*_ | *θ*), where *θ* is a vector of parameters that we will estimate. We collect the noisy data into a matrix denoted by 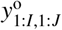, where the subscript makes it clear that the data consists of a time series of *I* observations with *J* measurements at each time point. Similarly, we denote the process model solution by *y*_1:*I*,1:*J*_(*θ*) for the matrix of predictions at the discrete time points and by *y*(*θ*) for the full (continuous) model trajectory over the time interval of interest [29].

We estimate parameters in the process models by assuming that the observed data are noisy versions of the model solutions of the form

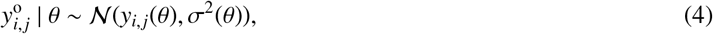

where *σ*^2^ is a constant variance that we also estimate from the data. In this work we take the simplest, most fundamental approach by assuming all measurements are subjected to the same noise model with the same constant variance. Our parameter vector, *θ*, is extended to include this constant variance and hence we write *σ*^2^(*θ*) for the variance component of *θ*. Regardless of whether we work with Equation (1) or (2)–(3), we make the assumption that the observation error is additive and normally distributed with zero mean and constant variance *σ*^2^ [42]. Different error models could be used within our likelihood-based framework if supported by the available data, for example working with a time-dependent variance if appropriate [4].

### 3.2. Parameter estimation

Combining our assumptions about process and noise models means that working with Equation (1) involves estimating four parameters, *θ* = (*λ, K, S* (0), *σ*), whereas working with Equations (2)–(3) requires the estimation of seven parameters *θ* = (*λ*_1_, *λ*_2_, *K, δ, C*_1_(0), *C*_2_(0), *σ*), where we make the simplifying assumption that the same noise model applies to all observations. Again, this assumption could be relaxed within our framework if the observed data justify doing so [4].

Taking a likelihood-based approach to parameter inference and uncertainty quantification, given a time series of observations together with our assumptions about the process and noise models, the log-likelihood function is

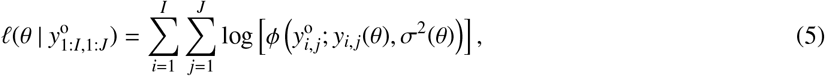

where *ϕ*(*x*; *µ, σ*^2^) denotes a Gaussian probability density function with mean *µ* and variance *σ*^2^. Maximum likelihood estimation (MLE) provides an estimate of *θ* that gives the *best* match to the data (in the sense of assigning the data the highest probability). The MLE is given by

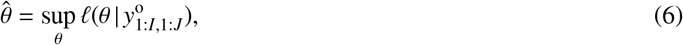

subject to bound constraints. The procedure for estimating 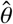 involves numerical maximisation of the log-likelihood, which can be achieved using many different algorithms. In this work, we find that a local optimisation routine from the open-source `NLopt` optimisation package in Julia performs well [21]. In particular, we use the Nelder-Mead optimisation routine within the `NLopt` with the default stopping criteria.

### 3.3. Practical identifiability analysis and profile predictions

We use a profile likelihood-based approach to explore practical identifiability by working with a normalised (relative) log-likelihood function

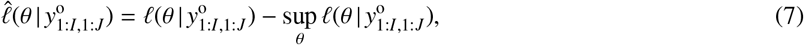

which we consider as a function of *θ* for a fixed set of data 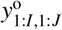. Normalising the log-likelihood ensures that 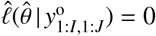.

#### 3.3.1. Profile likelihood for interest parameters

When the full parameter *θ* can be partitioned into an interest parameter *ψ* and nuisance parameter *ω*, where these may be vector valued in general, we write *θ* = (*ψ, ω*). More generally, an interest parameter can be defined as any function of the full parameter vector, *ψ* = *ψ*(*θ*); the nuisance parameter is then the implied complement, allowing the full parameter vector to be reconstructed. For a set of data, 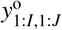, the profile log-likelihood for the interest parameter *ψ* given the partition (*ψ, ω*) is defined as [11, 31]

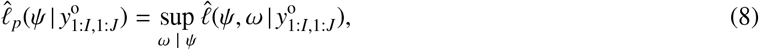

which indicates that *ω* is optimised out for each fixed value of *ψ*. This implicitly gives a function *ω*\*(***ψ***) of optimal values of *ω* for each value of *ψ*, and defines a curve (or, for non-scalar interest parameters, hypersurface) with points (*ψ, ω*\* (*ψ*)) in parameter space. In the case of an interest parameter defined by a function of the full parameter, the profile (or induced) log-likelihood is defined by the constrained optimisation problem [7, 10, 30]

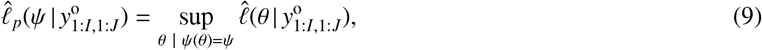

in which the remaining degrees of freedom (implicit nuisance parameters) in *θ*, after fixing *ψ*, are optimised out. Note that if we consider *ψ* to be a scalar interest parameter then (*ψ, ω*\* (*ψ*)) defines a univariate function that we can visualise as a curve. In contrast, if *ψ* is a pair of interest parameters then (*ψ, ω*\* (*ψ*)) is a function of two variables that is often called a bivariate profile that we can visualise as a heat map or contour plot [40].

To demonstrate, consider working with Equations (2)–(3) where *θ* = (*λ*_1_, *λ*_2_, *K, δ, C*_1_(0), *C*_2_(0), *σ*). If we wish to profile *λ*_2_ we have *ψ*(*θ*) = *λ*_2_ and *ω*(*θ*) = (*λ*_1_, *K, δ, C*_1_(0), *C*_2_(0), *σ*) so that

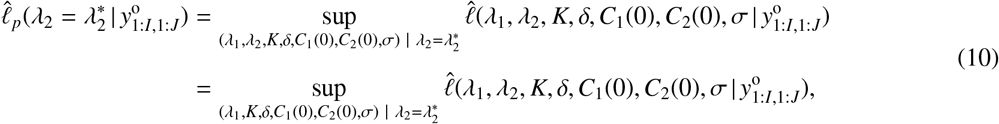

noting that we could write 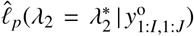 equivalently as 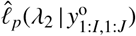. Here we chose the former notation to make it clear that we construct the univariate profile of *λ*_2_ by maximising the log-likelihood function at fixed values of 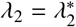.

In all cases, we implement this numerical optimisation using the same Nelder-Mead routine in NLopt that we use to estimate the MLE, 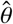 [21]. Our approach to computing the profile is very straightforward; we simply define a single uniform-spaced mesh across an interval in the interest parameter that contains the MLE approximately in the centre of the interval. Since the computations are relatively fast we can rapidly assess whether our choice of the width of the interval is sufficient. If, for example, we find that the interval leads to a profile that does not intersect the appropriate threshold we simple re-compute the profile on a wider interval. For each profile in this work we use a mesh of 40 uniformly spaced mesh points, and for each numerical optimisation calculation we simply use the MLE as the initial estimate for the iterative solver. The choice of working with a mesh of 40 uniformly spaced points can be varied easily, and this depends on the shape of the resulting profile. If the profile gradually varies across the interval of interest, as we find in this study, 40 mesh points appears to be sufficient to describe the shape of the profile; however if the profiles are relatively rapidly varying then we can easily repeat the calculations using more mesh points to resolve the shape of the profile. With these profiles, log-likelihood-based confidence intervals can be defined from the profile log-likelihood by an asymptotic approximation in terms of the chi-squared distribution that holds for sufficiently regular problems [31, 36]. For example, 95% confidence intervals for a univariate (scalar) interest parameter correspond to a threshold profile log-likelihood value of −1.92 [36]. Other confidence intervals can be estimated by changing the threshold value accordingly.

#### 3.3.2. Predictive profile likelihood and parameter-wise profile predictions

As discussed in [22, 29, 51], profile likelihoods for *predictive* quantities that are a deterministic function of the full parameter *θ*, such as model mean trajectories *y*(*θ*), are defined in the same way as for any other function of the full parameter (described in Section 3.3.1). For example, the model mean trajectory has an associated profile likelihood

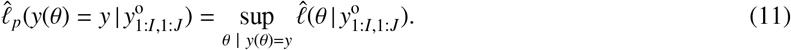

This defines the profile likelihood value for a prediction to be equal to the maximum likelihood value across all parameters giving that prediction. In principle, this gives the profile prediction for the full, continuous, model trajectory. However, this can be difficult to implement in practice and the literature typically focuses on a single-time prediction [22, 51].

In the present work, we are interested in some measure of the dependence of predictions on a given interest (target) parameter. However, as discussed in [29], given a partition *θ* = (*ψ, ω*) and a function *q*(*θ*) of the full parameter, in general the quantity *q*(*ψ*) is not well defined, unless *q* is independent of *ω*. To overcome this, we consider *parameter-wise profile predictions* [29], rather than the standard predictive profile likelihood discussed above. These consider the dependence of a predictive function *q*(*ψ, ω*) of the full parameter on the interest parameter *ψ* via its value along the corresponding profile curve, i.e. by considering *q*(*ψ, ω*\* (*ψ*)) where *ω*\* (*ψ*) is the optimal value of the nuisance parameter for a given value of the interest parameter. As discussed in [29], the transformation of profile confidence intervals for *ψ* into confidence intervals for the predictive quantity of interest *q*(*ψ, ω*) will, in general, only be approximate due to the additional dependence on nuisance parameters. However, we can use these parameter-wise intervals as an intuitive diagnostic tool that can reveal the influence of an interest parameter on predictions, as well as explore their coverage properties via simulation. In contrast, a standard predictive profile cannot reveal the individual influence of particular parameters as all parameters are simultaneously optimised over.

Formally, the associated profile likelihood for *q*(*ψ, ω*\* (*ψ*)) is defined by [29]:

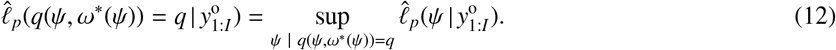

We compute this quantity by first evaluating the predictive quantity *q* along the profile curve (*ψ, ω*\* (*ψ*)) and assigning it tentative likelihood values equal to the associated profile likelihood value of (*ψ, ω*\* (*ψ*)). Then *q*(*ψ, ω*\* (*ψ*)) is assigned a single profile likelihood value equal to the maximum profile likelihood value over compatible *ψ* values (the ‘sup’ step). If *q*(*ψ, ω*\* (*ψ*)) is 1-1 in *ψ* then this value is simply the profile likelihood value of *ψ*, but it is well-defined regardless. This is analogous to constructing a profile for an interest predictive quantity *q* using a standard likelihood in *θ* but now starting from a profile likelihood for *ψ*. This makes it much easier to implement computationally – for a one-dimensional interest parameter we only need to evaluate the predictive quantity along a one-dimensional curve embedded in parameter space. For more on this definition, see [29]. Since we are evaluating *q* along the parameter profile curve (*ψ, ω*\* (*ψ*)), the likelihood thresholds defining approximate confidence sets for the predictive quantities are the same as for the parameters.

The above process allows us to construct an approximate interval for the predictive quantity assuming the uncertainty is driven primarily by the parameter of interest. As a sensitivity tool, this analysis illustrates which aspects of a prediction depend on the interest parameter. For example, the early part of the mean trajectory may have a stronger dependence on a given parameter than the later part of the trajectory. This feature of this tool is desirable from the point of view of understanding the impacts of individual parameters on predictions, but also means the individual intervals will in general be expected to have lower coverage for the prediction than the associated profile parameter interval (due to neglecting additional dependence on/uncertainty in the nuisance parameters). To overcome this issue to some degree, when interested in overall uncertainty we can combine the individual intervals to give an approximate overall interval. In particular, given a collection of individual intervals for the same quantity based on different interest parameters, more conservative confidence intervals for that quantity can be constructed by taking the union over all intervals. As an illustration, given two intervals (or sets) 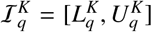 and 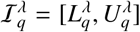 for a quantity *q* based on profiles for *K* and *λ*, respectively, we can form an interval (or set) 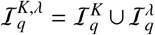 which has coverage at least as great as the individual intervals. In the case where the two intervals overlap, we have 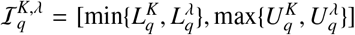. As with the individual intervals, the precise coverage properties of these combined intervals are difficult to establish formally without direct simulation, but such union intervals can provide an approximate picture of the overall variation that is efficient to compute and intuitive to understand.

With these definitions, we will now compute various univariate profile likelihoods, parameter-wise prediction intervals and the union of the parameter-wise prediction intervals for both Equation (1) and Equations (2)–(3).

## 4. Results

### 4.1. Single species model

Figure 2(a) compares the solution of Equation (1) evaluated at 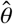 with the data, while in (b)–(c) univariate profiles are shown for each parameter. In each case, the univariate profiles are regularly shaped with clearly defined peaks, indicating that the model is practically identifiable [42]. Asymptotic confidence intervals for each parameter are determined by interpolating the profiles at the threshold of *ℓ* = −1.92 and reported in the caption of Figure 2.

**Figure 2:**
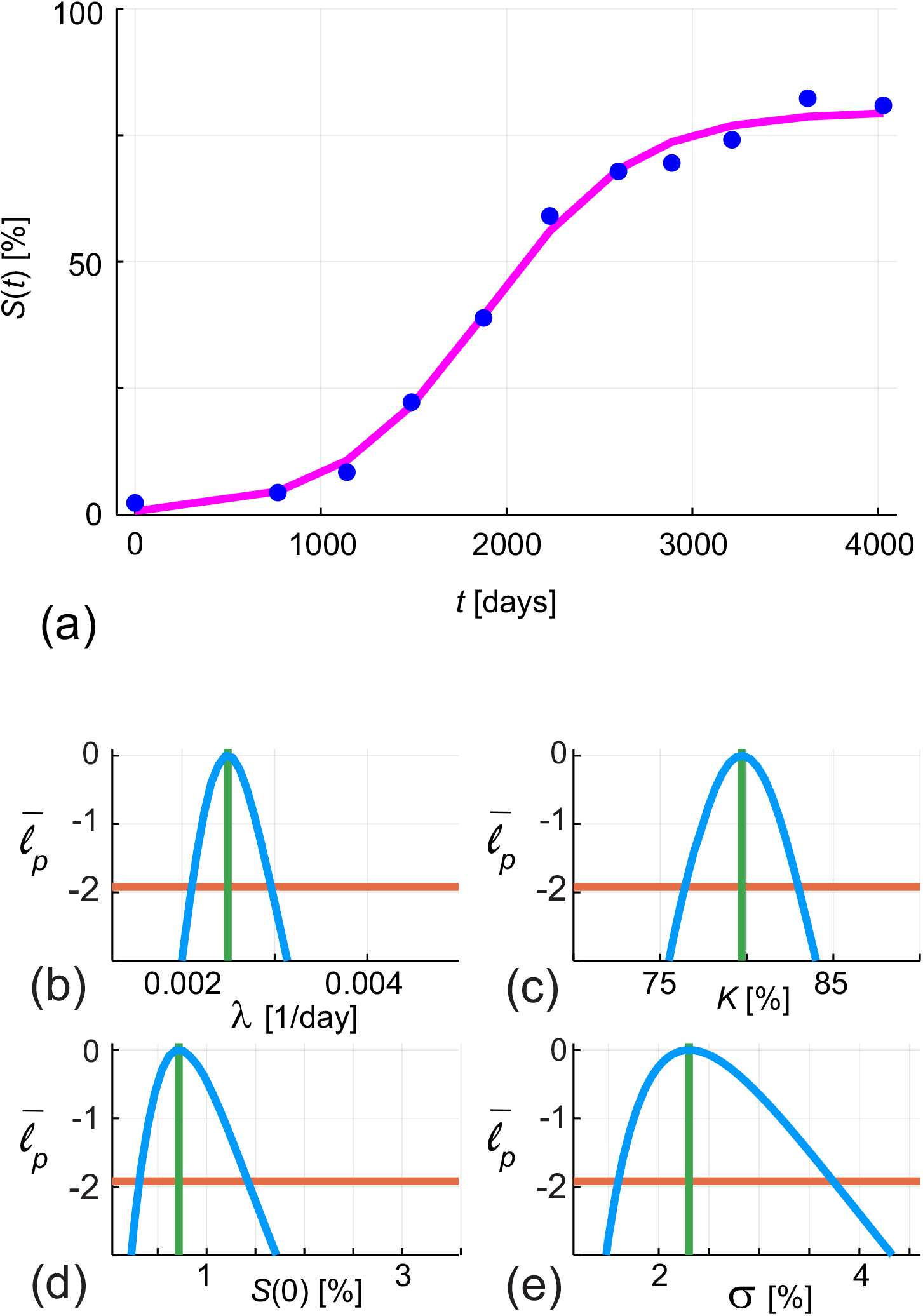
(a) Comparison of data (blue discs) and the solution of Equation (1) evaluated at the MLE, 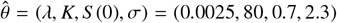 (pink line). (b) Univariate profile for *λ* gives 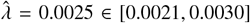. (c) Univariate profile for *K* gives 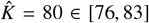. (d) Univariate profile for *S* (0) gives 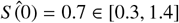. (e) Univariate profile for *σ* gives 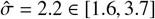. Each profile is shown in solid blue; the MLE as a vertical green line, and the asymptotic log-likelihood threshold as a horizontal orange line.

The profiles in Figure 2 provide important quantitative information about the extent to which the quality and quantity of data identifies the parameters in the mathematical model. For example, the MLE for the carrying capacity density gives 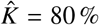, but the profiles allow us to identify the credible interval for this estimate, which tells us that the 95% credible interval is 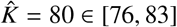 in this case, which provides us with a clear quantitative measure of the uncertainty in our estimate.

With the univariate profiles in Figure 2 we construct parameter-wise profile predictions shown in Figure 3 for *λ, K* and *S* (0), respectively. The width of the prediction interval for the parameter is correlated with the strength of the parameter in influencing the prediction of *S* (*t*), and the trends we see are intuitively reasonable. For example, the prediction interval associated with *K* is relatively narrow at early time, but widens considerably at later times. This is entirely reasonable since we know that *S* (*t*) → *K* as *t* → ∞ in the model. Similarly, the prediction interval associated with *S* (0) leads to a relatively wide prediction interval at early time, which is reasonable since the value of *S* (0) influences the early time dynamics, but the long-time solution of (1) is independent of *S* (0), since *S* (*t*) → *K* as *t* → ∞ regardless of *S* (0). Note that we do not construct a prediction interval for *σ* since this is a measure of the noise in the data and the solution of (1) is completely independent of *σ* [29].

**Figure 3:**
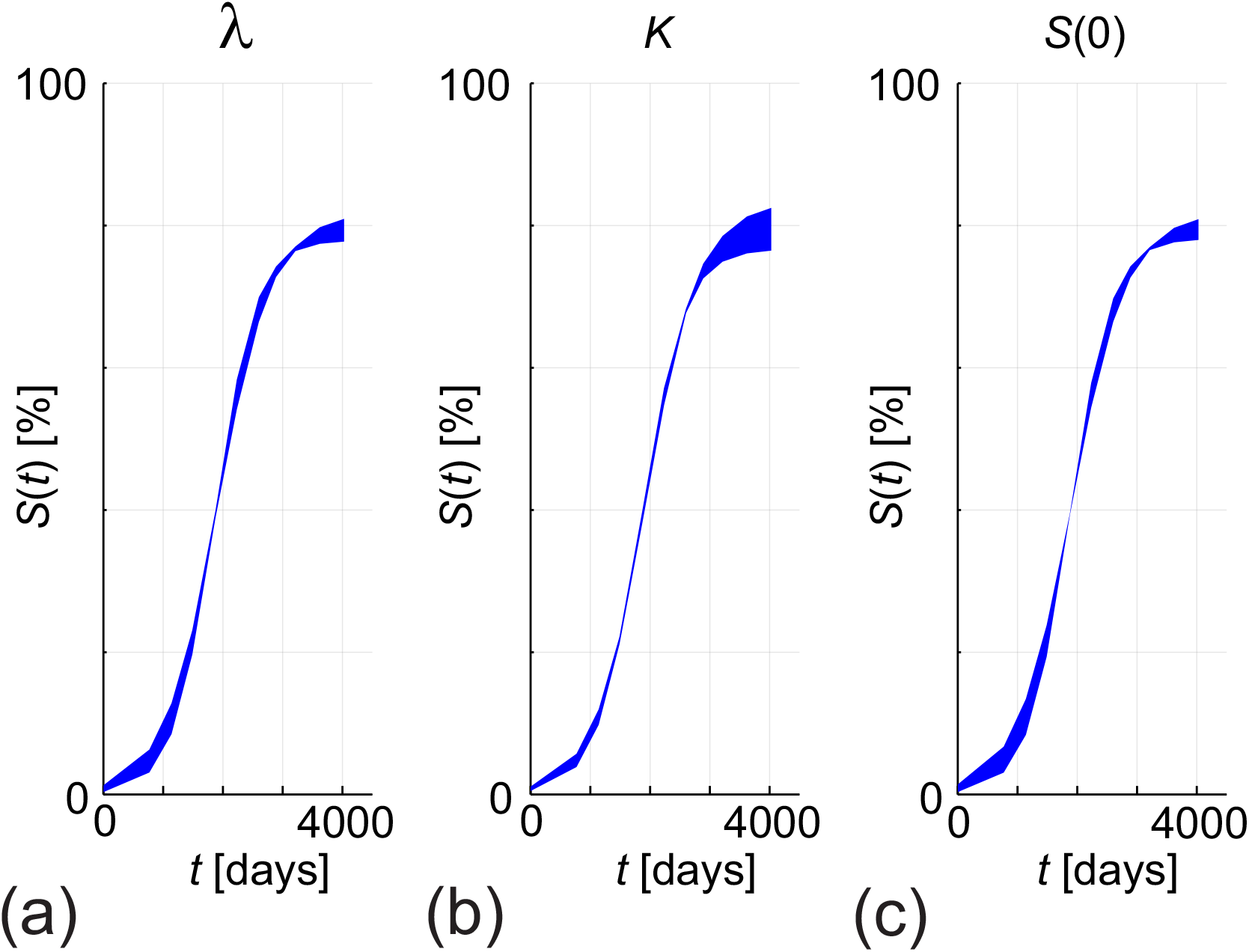
Parameter-wise profile predictions for modelling total hard coral coverage with Equation (1). Results in (a)–(c) show the 95% parameter-wise profile predictions for *λ, K* and *S* (0), respectively.

### 4.2. Two species model

Figure 4(a)–(b) compares the solution of Equations (2)–(3) evaluated at 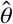 with the data Figure 4(c) compares *S* (*t*) = *C*_1_(*t*) + *C*_2_(*t*) with the data describing the evolution of the total hard corals. Univariate profiles are given for each parameter in Figure 4(d)–(j); again, in each case, the profiles are regularly-shaped with clearly defined peaks, suggesting that this more complicated model is also practically identifiable with the data. Asymptotic confidence intervals for each parameter are determined by interpolating the profiles at the threshold of *ℓ* = −1.92 and reported in the caption of Figure 4.

**Figure 4:**
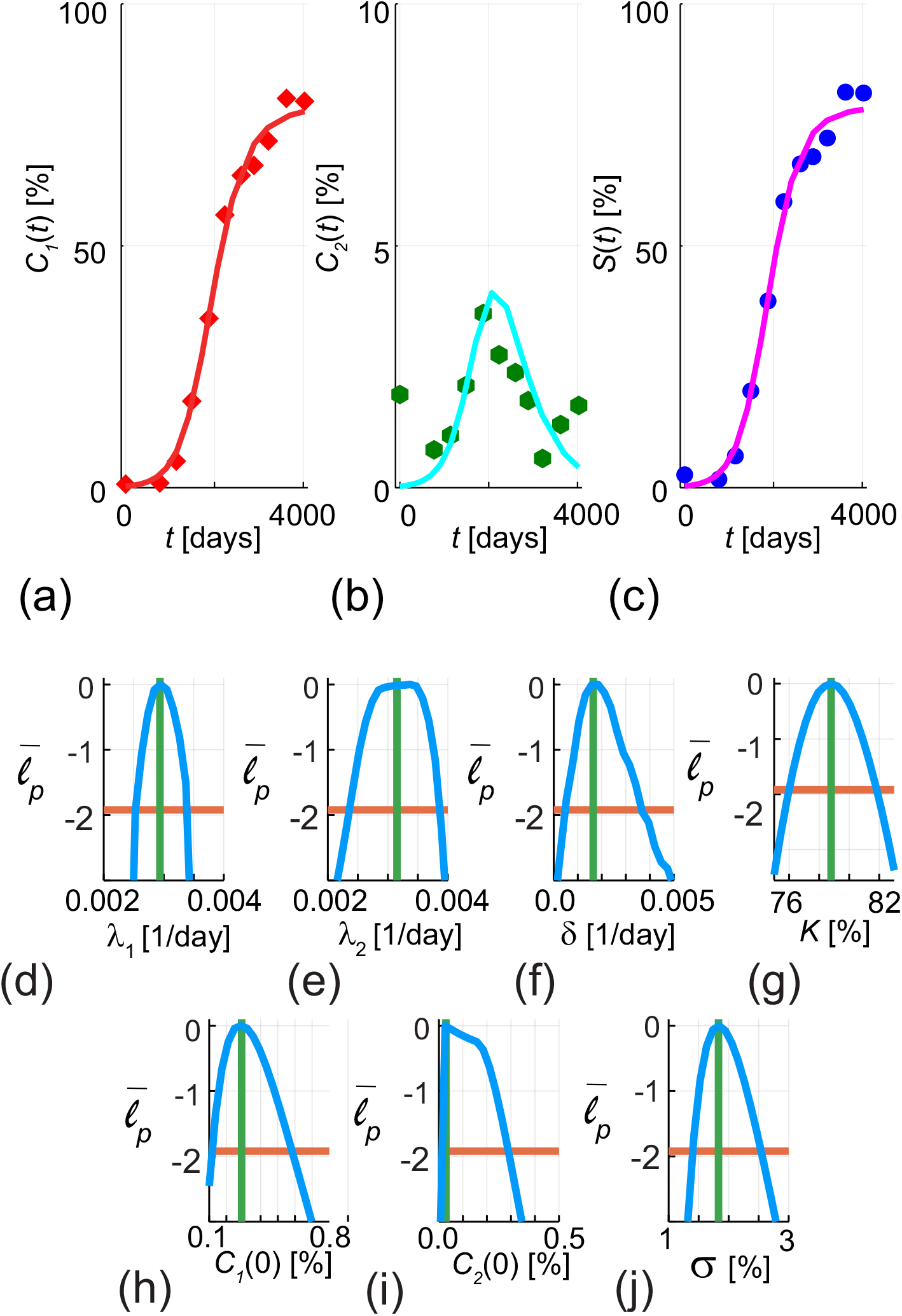
(a)–(b) Comparison of data and the solution of Equations (2)–(3) for *C*_1_(*t*) and *C*_2_(*t*), respectively, evaluated at the MLE, 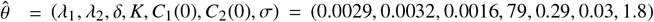. Results in (c) compare data and the solution of (2)–(3) in terms of the total hard coral coverage, *S* (*t*), evaluated at 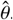. Results in (a) compare the data (red diamonds) with the MLE solution (solid red curve), results in (b) compare the data (green dots) with the MLE solution (solid cyan curve). results in (c) compare the data (blue dots) with the MLE solution (solid pink curve). (d) Univariate profile for *λ*_1_ gives 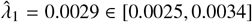. (e) Univariate profile for *λ*_2_ gives 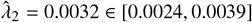. (f) Univariate profile for *δ* gives 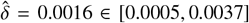. (g) Univariate profile for *K* gives 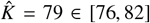. (h) Univariate profile for *C*_1_(0) gives 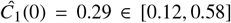. (i) Univariate profile for *C*_2_(0) gives 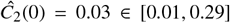. (j) Univariate profile for *σ* gives 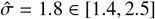. Note that *C*_2_(*t*) is plotted on a different vertical scale than *C*_1_(*t*) or *S* (*t*). Each profile in (d)–(j) is shown in solid blue; the MLE as a vertical green line, and the asymptotic log-likelihood threshold as a horizontal orange line.

With the univariate profiles in Figure 4, we construct parameter-wise profile predictions, shown in Figure 5, for each parameter in Equations (2)–(3). In this case we can plot prediction intervals for *C*_1_(*t*), *C*_2_(*t*) and for *S* (*t*) = *C*_1_(*t*) + *C*_2_(*t*) as a function of each parameter in the process model. Again, these results are intuitively reasonable since we see, for example, that the prediction interval for *λ*_2_ has a more significant influence on the prediction interval for *C*_2_(*t*) than *C*_1_(*t*), and that the prediction interval associated with *K* has an influence on the late-time behaviour of *C*_1_(*t*), *C*_2_(*t*) and *S* (*t*) and a negligible influence on the early time prediction intervals for any of these three quantities.

**Figure 5:**
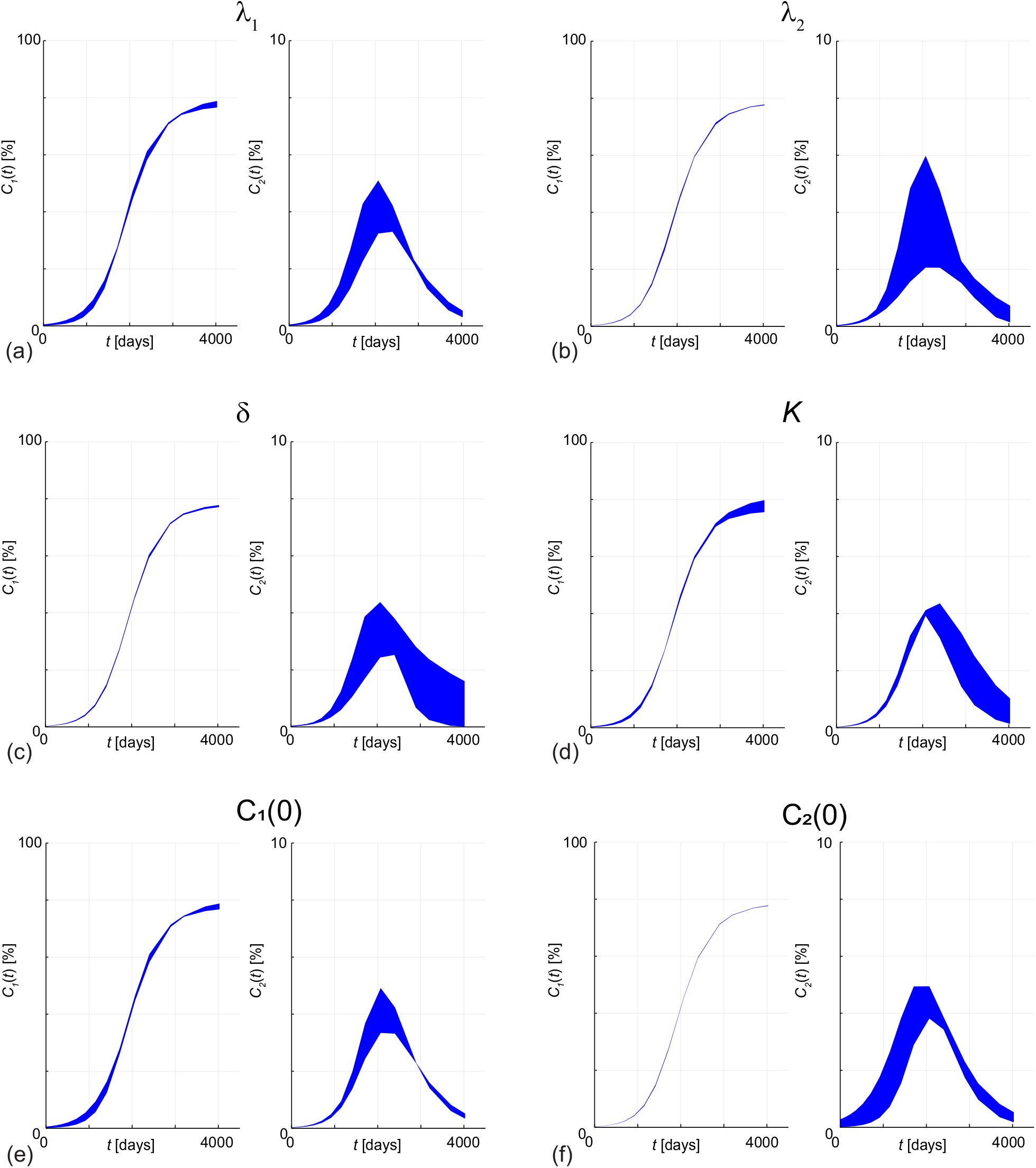
Parameter-wise profile predictions for modelling coral coverage with Equations (2)–(3) in terms of *C*_1_(*t*), *C*_2_(*t*) and *S* (*t*) = *C*_1_(*t*) + *C*_2_(*t*). Results in (a)–(f) show the 95% parameter-wise profile predictions for *λ*_1_, *λ*_2_, *δ, K, C*_1_(0) and *C*_2_(0), respectively. Note that *C*_2_(*t*) is plotted on a different vertical scale than *C*_1_(*t*) or *S* (*t*).

### 4.3. Comparing the single- and two species modelling approaches

With our profile likelihood results constructed for Equation (1) in Figure 2 and for Equations (2)–(3) in Figure 4 we are able to make several comparisons between the two modelling approaches. For example, we can simply compare parameter point and interval estimates. To illustrate, for Equation (1) we have a point estimate 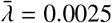 and interval estimate [0.0021, 0.0030], whereas for Equations (2)–(3) we have point estimates 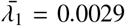 and 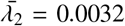 and interval estimates [0.0025, 0.0034] and [0.0024, 0.0039] for *λ*_1_ and *λ*_2_, respectively. This suggests that the growth rates in the two models are largely statistically indistinguishable, with slightly greater uncertainty in *λ*_2_ in the coupled model.

Comparing the uncertainty, as measured by the widths of confidence intervals, associated with our more complex model can inform our opinion about which modelling approach is preferred [42]. For example, with the two-species model we have 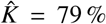 which is close to the MLE for the single species model, but we can also quantitatively compare the widths of the confidence intervals. Here for the two species model we have 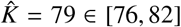, whereas for the single species model we have 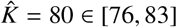, from which we can conclude that our estimate of the carrying capacity density, and the uncertainty in that estimate is not very different between the two modelling approaches despite significant differences between the two modelling approaches. However, instead of just comparing models via uncertainties in parameter estimates, it is perhaps more natural in many applications to convert these to associated predictive intervals for a predictive quantity of interest such as *S* (*t*), given in Figure 6. This figure shows the union of the various parameter-wise prediction intervals for each model. Here we see, despite the differences in both models, in this application the prediction intervals are not strikingly different. Thus the overall uncertainty in one of the key quantities of interest is largely the same for both models.

**Figure 6:**
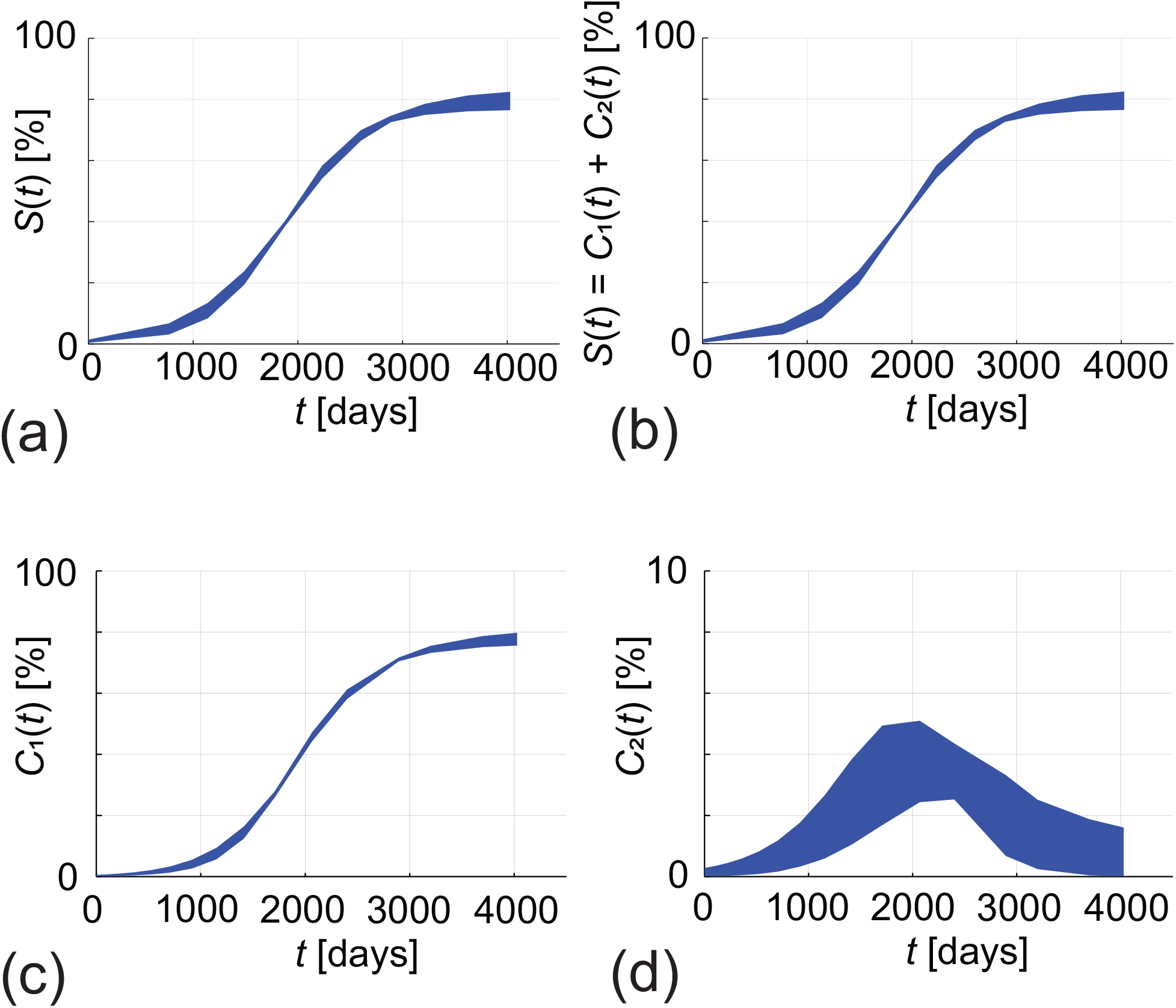
(a)–(b) gives the union of the parameter-wise profile predictions using (1) and (2)–(3) for *S* (*t*), respectively. The prediction intervals in (a)–(b) are formed by taking the union of the prediction intervals in Figure 3 and Figure 5, respectively. The prediction intervals in (c)–(d) are formed by taking the union of the prediction intervals in Figure 5 for *C*_1_(*t*) and *C*_2_(*t*), respectively. Note that *C*_2_(*t*) is plotted on a different vertical scale than *C*_1_(*t*) or *S* (*t*).

There are many ways that we can interpret these results. If, for example, the main interest in modelling the coral regrowth is simply to predict the total coral coverage, then our comparisons in Figure 6 suggest that working with Equation (1) may be preferable since the prediction interval for the quantity of interest is not very different between the simpler model and the more complicated model, and of course working with the simpler model has pragmatic advantages. Alternatively, if the aim of the modelling exercise is to use a mathematical model to highlight the distinction between the regrowth of *Acroporidae* and other hard corals in the community, then the comparison in Figure 6 could be used to indicate that working with Equations (2)–(3) is entirely reasonable. For example, despite the relative complexity of the coupled model, working with the coupled model does not lead to significantly wider prediction intervals for the quantity in common with the simple model, although it does provide intervals for the separate components.

As we stated in Section 4.2, parameter-wise prediction intervals are approximate in the sense that the transformation of confidence intervals for *ψ* into confidence intervals for the predictive quantity is not exact owing to non-trivial dependencies between nuisance parameters. For simple models, such as Equation (1), it is straightforward to explore the impact of this approximation by constructing prediction intervals from the full likelihood and comparing these with the union of parameter-wise prediction intervals. Additional results in the Appendix for Equation (1) show the union of parameter-wise intervals does indeed lead to smaller uncertainty estimates than evaluating the full likelihood directly, although the differences are relatively small in this case and the increase in computational effort for the full likelihood is significant.

In addition to comparing the single species and two species models through the lens of profile likelihoods and parameter wise profile predictions, it is also instructive to compare our approach with more traditional information criteria based analysis [1]. One standard way to compare models is using the Akaike Information Criteria (AIC) [1]

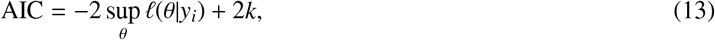

where *k* is the dimension of *θ*. Given a set of candidate models describing the same data, the preferred model is the one with the minimum AIC [1]. This kind of analysis rewards goodness of fit (as measured by the log-likelihood function) and penalises model complexity (as measured by the dimensionality of *θ*). This approach is very easy to implement and very popular and enables us to quantitatively compare two different models of the same data by ranking them in terms of AIC [48]. However, while information criteria-based measures describe the tradeoff between the goodness-of-fit and the dimensionality of the parameter space, estimates of AIC do not provide insight into the shape of the likelihood function, thereby providing no insight into the practical identifiability of the parameters. This is important because, in general, the curvature of the likelihood is related to inferential precision and so our approach provides additional insight by combining information about the goodness of fit through of the model (indicated by the MLE), the shape of the likelihood function (indicated by the width of the profile), as well as the width of the prediction intervals.

## 5. Conclusions and Outlook

In this work we provide a straightforward means of comparing mathematical models of population dynamics, in particular we use a case study describing coral reef regrowth after some disturbance. Our focus is to compare models and data using two different approaches. First, we study the time evolution of the total hard coral cover, *S* (*t*). Second, we divide up the total population of hard corals into two groups, the dominant hard coral group *Acroporidae, C*_1_(*t*), and all other hard corals lumped together, *C*_2_(*t*), such that *S* (*t*) = *C*_1_(*t*) + *C*_2_(*t*). Our modelling philosophy is always to use a minimal model where possible, so in the first approach where we make no distinction between the different types of hard corals, we model the total hard coral cover, *S* (*t*), using a classic logistic growth model. In the second approach, where we explicitly model the regrowth of both types of hard corals, we use a system of coupled ODEs. This system involves logistic growth-like terms for each type of hard coral, together with a competition term describing how the dominant hard coral group competes with the remaining hard corals. Both of these ODE-based models are structurally identifiable, and univariate profile likelihood analysis confirms that, with our data, both the simple single species logistic model and the more complicated two-species models are practically identifiable. This is an important result since it is unclear when we move from the simple four parameter logistic model to the more complicated seven parameter coupled model whether the parameters in the more complicated model are identifiable. To provide more insight into the performance of both modelling approaches we construct and compare parameter-wise prediction intervals for both approaches. Our approach to constructing parameter-wise prediction intervals is novel and computationally straightforward, and reveals how each parameter impacts the predictions of the mathematical models. Understanding how model prediction is impacted by variability in each parameter is insightful since this gives us the ability to understand how uncertainty in each parameter contributes to the overall variability in the quantity of interest. Our results show that both modelling approaches lead to similar prediction outcomes. Therefore, choosing which modelling approach suits our needs depends strongly upon the aim of the modelling study; if the aim of the modelling study is to understand the total hard coral cover then it is entirely reasonable to work with the simpler single-species logistic model for pragmatic reasons. In contrast, if the aim of the modelling study is to understand interactions of different coral groups within the total population of hard corals, then working with the more complicated model does not lead to any parameter identifiability issues, nor is there any significant difference in the predictive capability of either modelling approach.

One of the aims of this work is to produce a fairly straightforward set of algorithms that can be used to guide model choice more broadly in the fields of mathematical biology and mathematical ecology. Model choice remains an on-going challenge in both mathematical biology and mathematical ecology, in part because data collection is not standardised and in part because there are many competing models available to describe similar phenomena [16, 37]. Within the context of modelling coral regrowth there are many other modelling approaches that one could take. For example, in this study we have compared working with total coral cover, *S* (*t*), with a more detailed approach where we divide hard corals into two groups: *C*_1_(*t*) and *C*_2_(*t*). Another approach would be to divide up the hard corals into more than two groups, giving *C*_*j*_(*t*) for *j* = 1, 2, 3, … *J*, and to then work with a system of *J* coupled ODEs to describe the evolution of the community of hard corals. In this work we focus on setting *J* = 2 because it is clear from the field data in Figure 1(c) that the total hard corals is dominated by *Acroporidae*, and so it is pragmatic to set *J* = 2 so we capture the evolution of *Acroporidae* and the evolution of all other hard coral types. In a different scenario, where no single type of hard coral dominates the total population, we might choose to work with *J* > 2, and in this case the tools presented here could be applied in a very similar way. Another extension would be to maintain working with *J* = 2 groups of hard coral in the more complicated mathematical model but to also include an additional competition term,

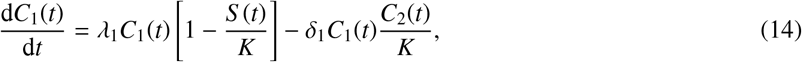

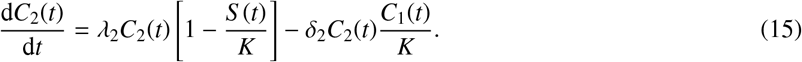

This extended model is identical to Equations (2)–(3) with the exception that now we explicitly include the possibility of the other hard corals competing with the *Acroporidae* through the parameter *δ*_1_, whereas in Equations (2)–(3) we implicitly assume that *δ*_1_ = 0 because there is no evidence of *C*_1_(*t*) decreasing with time in our data. This extended two-species model could be applied to our data using the same methodology and computational tools that we have developed. Of course, this particular extension is just one of many kinds of modelling extensions that we could analyse with our framework. Another extension would be to consider separate carrying capacity densities *K*_1_ and *K*_2_ for each coral type *C*_1_ and *C*_2_, respectively. While, in general, it would be reasonable to assume that different coral types might grow to reach different carrying capacity densities, here in our data we see that *C*_1_(*t*) dominates at late time with *C*_2_(*t*) → 0 for large *t* and so our data does not characterise late time maximum density behaviour for *C*_2_(*t*), and our previous experience in estimating late-time high-density parameters using early-time, low-density data leads to identifiability issues [47] so we do not pursue this extension here. Another feature of our modelling approach is that we focus on deterministic ODE models with a separate noise model. While this is an important class of widely-used models, other approaches such as working with stochastic process models are also of high interest for different applications [40] and an interesting extension of the current work would be to extend the current framework so that we can deal with stochastic process models. An intermediate approach would be to use a deterministic ODE model but correct the noise model to account for missing stochastic process model effects [41].

In addition to simply applying the tools developed here to different mathematical models, the current work lays the foundation for future theoretical developments. For example, results in the Appendix compare predictions for the simple single-species logistic model, Equation (1), obtained by taking the union of our parameter-wise prediction intervals with the prediction intervals obtained by evaluating the likelihood function directly. While working with a uniform discretisation of the parameter space to evaluate the full likelihood is reasonable for a simple model, this approach rapidly becomes computationally intractable for more complicated models like Equations (2)–(3). This is precisely the scenario where our parameter-wise prediction interval approach is advantageous. Nonetheless, our results for the simple model indicate that the union of the parameter-wise prediction intervals leads to slightly narrowed prediction intervals relative to evaluating the likelihood directly. This observation suggests further investigation is warranted and one approach would be to construct bivariate and higher order profiles, and take the union of a series of predictions made across these bivariate and higher order profiles [4]. While this approach would incur an increased computational cost over working with univariate profiles only, it would still be more efficient compared to evaluating the likelihood directly. We intend to explore the balance of computation time and accuracy in future work.

## Supporting information

Additional results and discussion

## Appendix: Comparing predictions from the full and profile likelihood

In this appendix we construct prediction intervals for *S* (*t*) using Equation (1), noting that for the simple model the log-likelihood function, 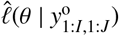 depends on just three parameters in the process model: *λ, K* and *S* (0). We evaluate the full log-likelihood function by defining a three-dimensional rectangular prism in *λ, K, S* (0) parameter space such that: 0.004 ≤ *λ* ≤ 0.001, 75 ≤ *K* ≤ 85 and 0 ≤ *S* (0) ≤ 1. We will justify the choice of these limits later. We then take a 30 × 30 × 30 uniform discretization of this rectangular prism and evaluate 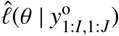 at each of the 27,000 points in the discretised parameter space. For those mesh points where 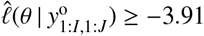 (corresponding to a 95% asymptotic confidence interval [36]) we plot the evolution of *S* (*t*) in Figure 7(a). Note that we chose the extent of the rectangular prism in *λ, K, S* (0) so that it completely enclosed the region where 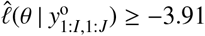.

**Figure 7:**
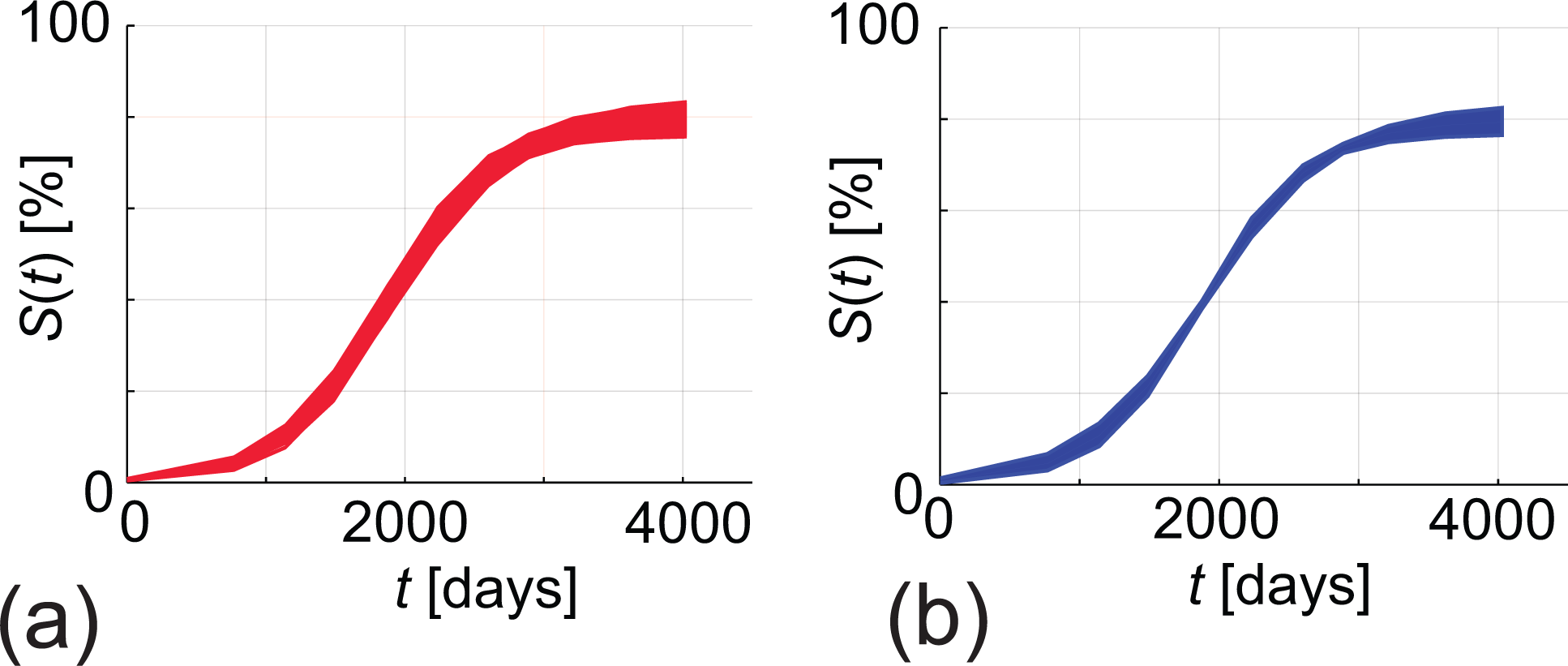
(a) prediction interval for the full likelihood (red curves). (b) approximate prediction interval constructed from profile likelihoods (blue curves).

The prediction intervals in Figure 7(a) for *S* (*t*) are generated by evaluating the log-likelihood directly. Visually comparing this collection of predictions with the union of the corresponding three parameter-wise prediction intervals in Figure 7(b) indicates that evaluating the full likelihood leads to similar, albeit slightly more conservative (broader), prediction intervals than those obtained from the parameter-wise approach. Therefore, in this case the parameter-wise intervals are slightly narrower than evaluating the full likelihood. However, the parameter-wise approach is both insightful (in terms of illustrating the impacts of each parameter) and computationally efficient since the computational effort required to compute the prediction intervals scales linearly with the dimension of the parameter space. For example here the prediction intervals are generated simply by considering 3 × 40 parameter combinations, whereas uniformly evaluating the log-likelihood function required 30 × 30 × 30 parameter combinations. While it is possible to use more sophisticated evaluation methods (e.g. Latin Hypercube Sampling) to work with the full likelihood function, this approach is likely infeasible for more complicated models where evaluating larger parameter spaces efficiently is a computational challenge. The parameter-wise approach, in contrast, remains computationally efficient even when the dimensionality of the full parameter space increases, since we rely on constructing the prediction intervals using a series of univariate profile likelihood functions only.

## Acknowledgements

MJS is supported by the Australian Research Council (DP200100177). SW and ES thank the Faculty of Science, Queensland University of Technology, for providing summer vacation research scholarships that supported this study. We thank three referees for helpful suggestions.

## Notes

https://github.com/ProfMJSimpson/profile_predictions

### Competing Interest Statement

The authors have declared no competing interest.

### Summary of Updates

Updated plots

