## Additional results and discussion for "Profile likelihood-based parameter and predictive interval analysis guides model choice for ecological population dynamics"

### S1. Introduction

In this Supplementary Material document we repeat all calculations from the main paper for a second data set, site 3 at Lady Musgrave Island. In brief, all of the general observations made in the main paper for site 1 carry across to our findings here. Furthermore, all results presented here can be replicated using the jupyter notebooks on GitHub

### S2. Results

#### *S2.1. Single species model*

Results in Figure S1 show the data for site 3, the solution of Equation (1) evaluated at the MLE and the univariate profiles for each parameter in the model. Details of the widths of the profiles are given in the caption and these results are all analogous to those results in Figure 3 in the main document for site 1.

Results in Figure S2 show the parameter-wise predictions for data for site 3 for each parameter in Equation (1), these results are all analogous to those results in Figure 2 in the main document for site 1.

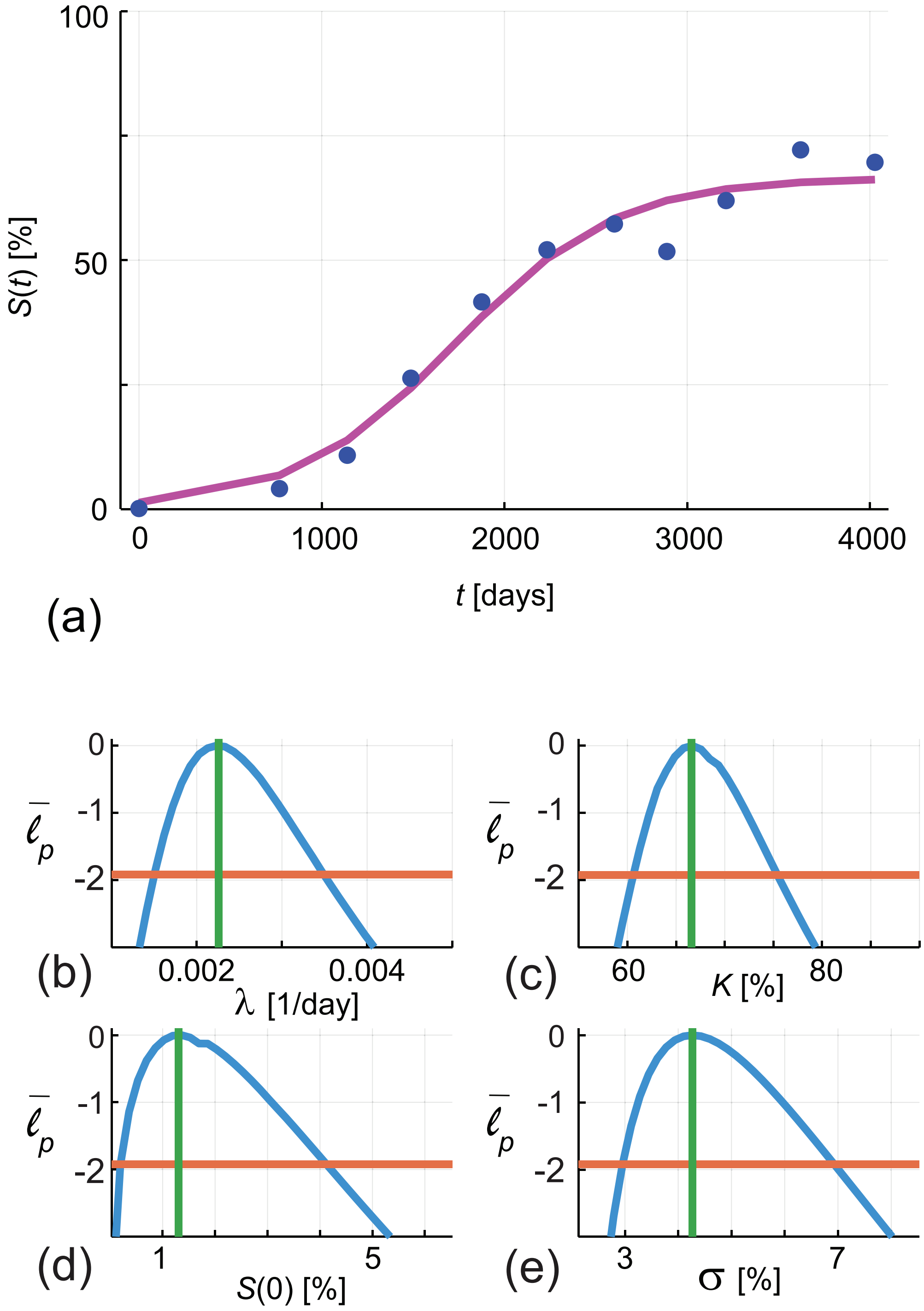


Figure S1: (a) Comparison of data (blue discs) and the solution of Equation (1) evaluated at the MLE, $\hat{\theta} = (\lambda, K, S(0), \sigma) = (0.0023, 67, 1.3, 4.3)$ (pink line). (b) Univariate profile for $\lambda$ gives $\hat{\lambda} = 0.0023 \in [0.0015, 0.0035]$. (c) Univariate profile for $K$ gives $\hat{K} = 67 \in [61, 75]$. (d) Univariate profile for $S(0)$ gives $S\hat{(0)} = 1.3 \in [0.2, 4.1]$. (e) Univariate profile for $\sigma$ gives $\hat{\sigma} = 4.3 \in [3.0, 6.9]$. Each profile is shown in solid blue; the MLE as a vertical green line, and the asymptotic log-likelihood threshold as a horizontal orange line.

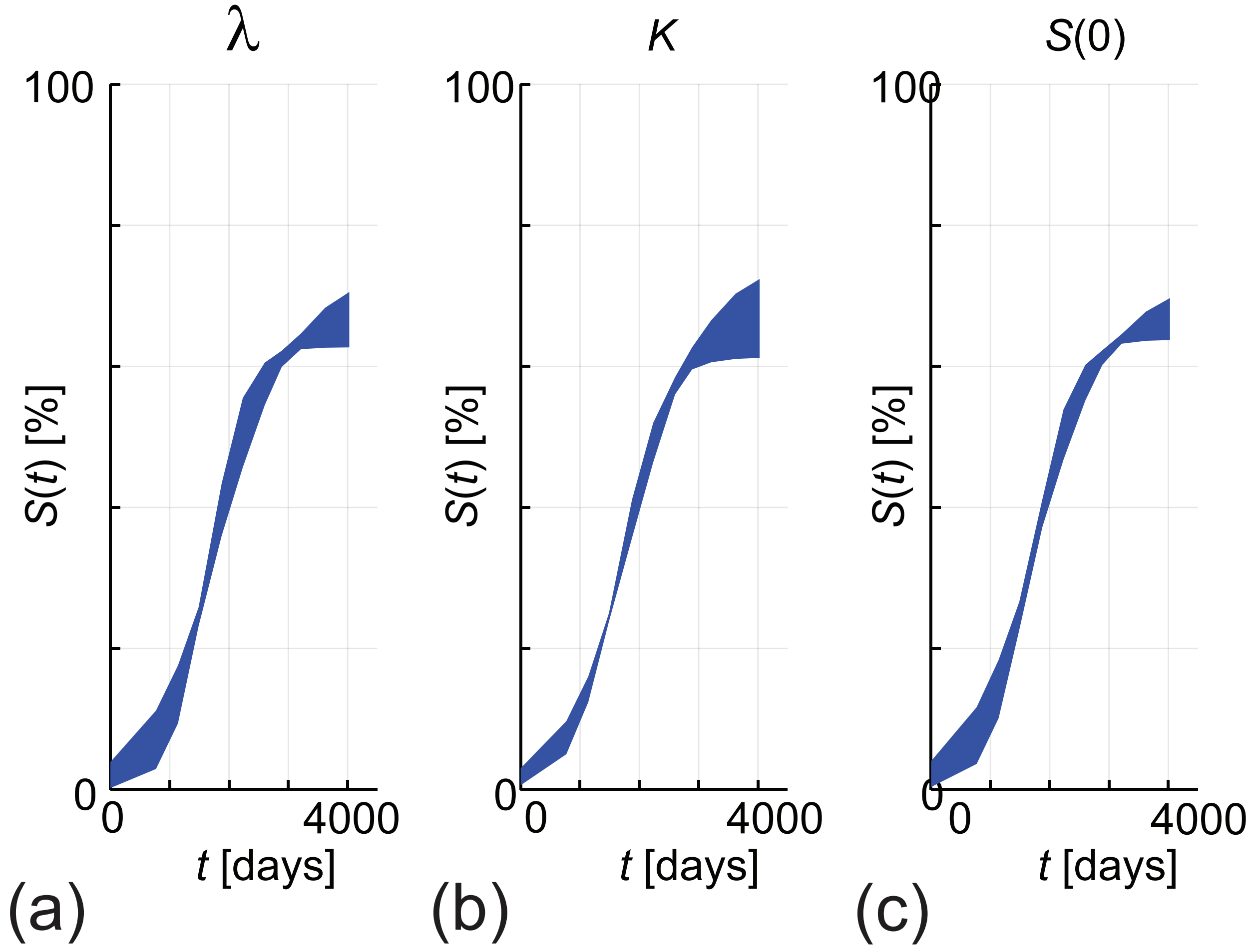


Figure S2: Parameter-wise profile predictions for modelling total hard coral coverage with Equation (1). Results in (a)–(c) show the 95% parameter-wise profile predictions for $\lambda$, $K$ and $S(0)$, respectively.

#### *S2.2. Two species model*

Results in Figure S3 show the data for site 3, the solution of Equations (2)–(3) evaluated at the MLE and the univariate profiles for each parameter in the model. Details of the widths of the profiles are given in the caption and these results are all analogous to those results in Figure 4 in the main document for site 1.

Results in Figure S4 show the parameter-wise predictions for data for site 3 for each parameter in Equations (2)–(3), these results are all analogous to those results in Figure 4 in the main document for site 1.

#### *S2.3. Comparing the single- and two species modelling approaches*

Figure S5 shows the union of the parameter-wise predictions using the single species model (1) and the two-species model (2)–(3) for data at site 3, these results are all analogous to those results in Figure 5 in the main document for site 1.

#### *S2.4. Comparing predictions from the full and profile likelihood*

Figure S6 compares the prediction intervals for the full likelihood and the union of prediction intervals associated with the single species model (1) for site 3. These results are all analogous to those results in Figure 7in the main document for site 1.

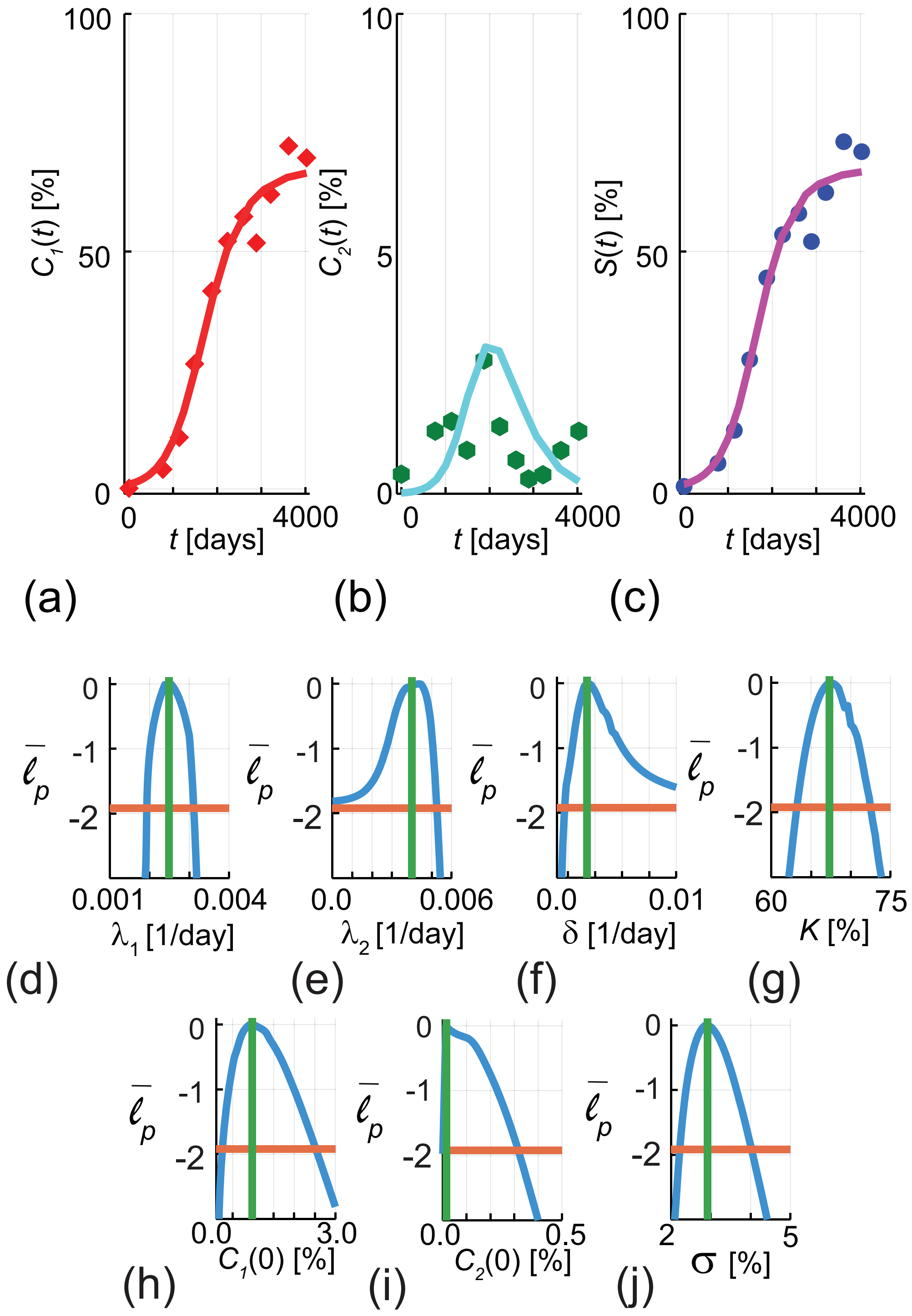


Figure S3: (a)–(b) Comparison of data and the solution of Equations (2)–(3) for $C_1(t)$ and $C_2(t)$, respectively, evaluated at the MLE, $\hat{\theta} = (\lambda_1, \lambda_2, \delta, K, C_1(0), C_2(0), \sigma) = (0.0025, 0.0040, 0.0018, 67, 0.97, 0.01, 2.9)$. Results in (c) compare data and the solution of Equations (1)–(2) in terms of the total hard coral coverage, $S(t)$, evaluated at $\hat{\theta}$. Results in (a) compare the data (red diamonds) with the MLE solution (solid red curve), results in (b) compare the data (green dots) with the MLE solution (solid cyan curve). results in (c) compare the data (blue dots) with the MLE solution (solid pink curve). (d) Univariate profile for $\lambda_1$ gives $\hat{\lambda}_1 = 0.0025 \in [0.0019, 0.0031]$. (e) Univariate profile for $\lambda_2$ gives $\hat{\lambda}_2 = 0.0040 \in [0.00, 0.0053]$. (f) Univariate profile for $\delta$ gives $\hat{\delta} = 0.0017 \in [0.00, 0.01]$. (g) Univariate profile for $K$ gives $\hat{K} = 67 \in [63, 72]$. (h) Univariate profile for $C_1(0)$ gives $\hat{C}_1(0) = 0.97 \in [0.25, 2.52]$. (i) Univariate profile for $C_2(0)$ gives $\hat{C}_2(0) = 0.01 \in [0.00, 0.32]$. (j) Univariate profile for $\sigma$ gives $\hat{\sigma} = 2.9 \in [2.2, 4.0]$. Note that $C_2(t)$ is plotted on a different vertical scale than $C_1(t)$ or $S(t)$. Each profile in (d)–(j) is shown in solid blue; the MLE as a vertical green line, and the asymptotic log-likelihood threshold as a horizontal orange line.

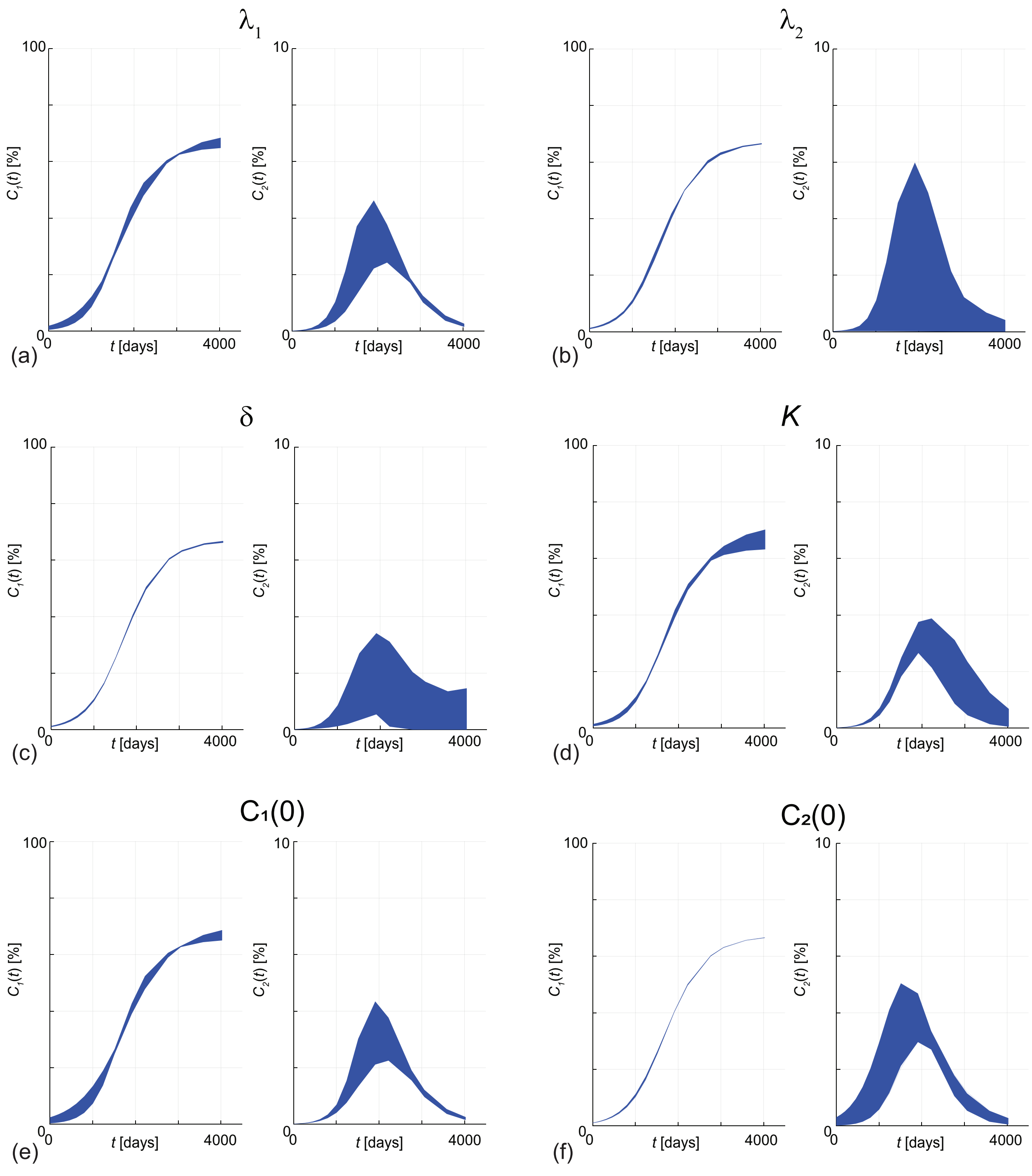


Figure S4: Parameter-wise profile predictions for modelling coral coverage with Equations (2)–(3) in terms of $C_1(t)$, $C_2(t)$ and $S(t) = C_1(t) + C_2(t)$. Results in (a)–(f) show the 95% parameter-wise profile predictions for $\lambda_1$, $\lambda_2$, $\delta$, $K$, $C_1(0)$ and $C_2(0)$, respectively. Note that $C_2(t)$ is plotted on a different vertical scale than $C_1(t)$ or $S(t)$.

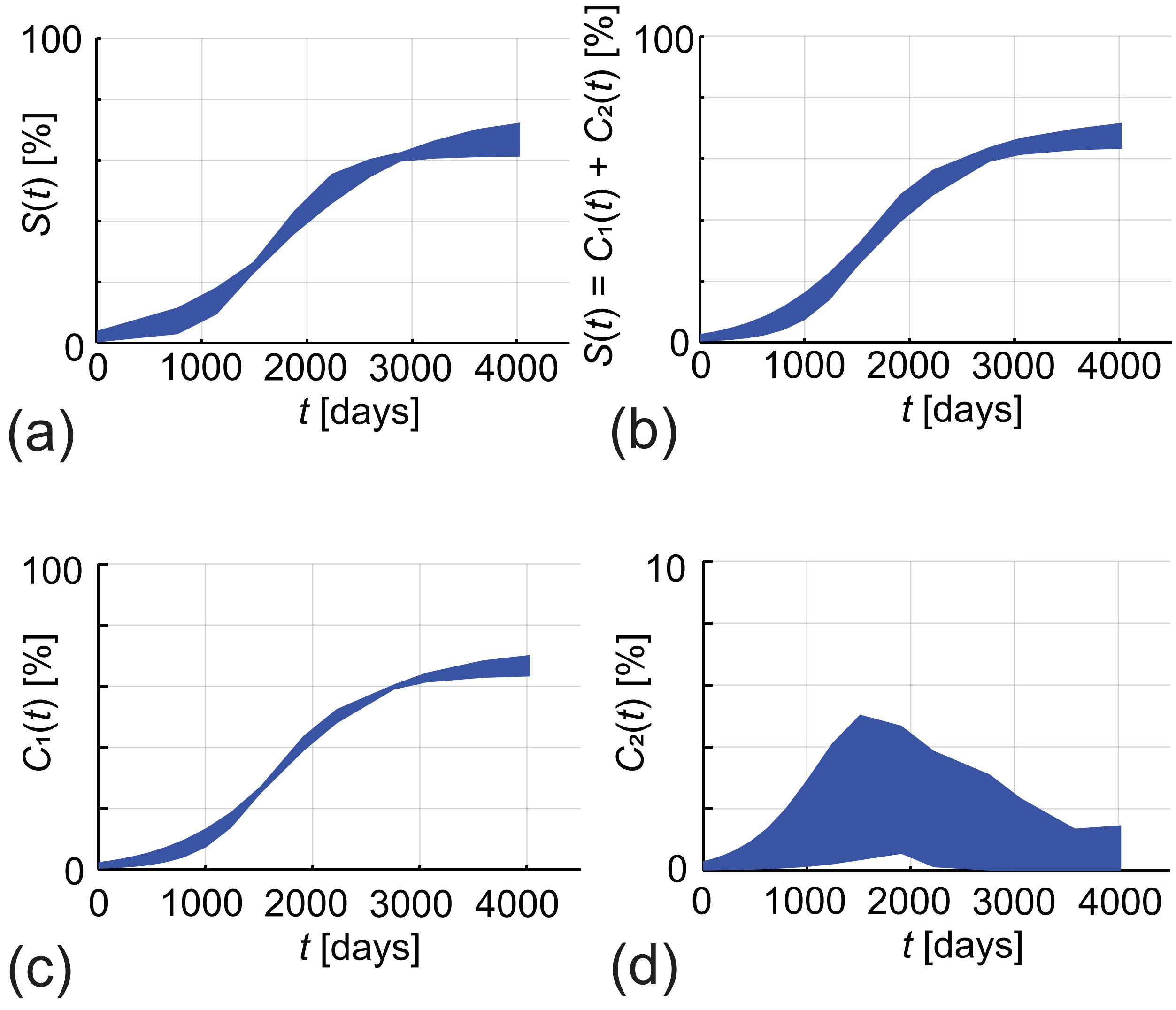


Figure S5: (a)–(b) gives the union of the parameter-wise profile predictions using Equation (1) and Equations (2)–(3) for $S(t)$, respectively. The prediction intervals in (a)–(b) are formed by taking the union of the prediction intervals in Figure S2 and Figure S4, respectively. The prediction intervals in (c)–(d) are formed by taking the union of the prediction intervals in Figure S4 for $C_1(t)$ and $C_2(t)$, respectively. Note that $C_2(t)$ is plotted on a different vertical scale than $C_1(t)$ or $S(t)$.

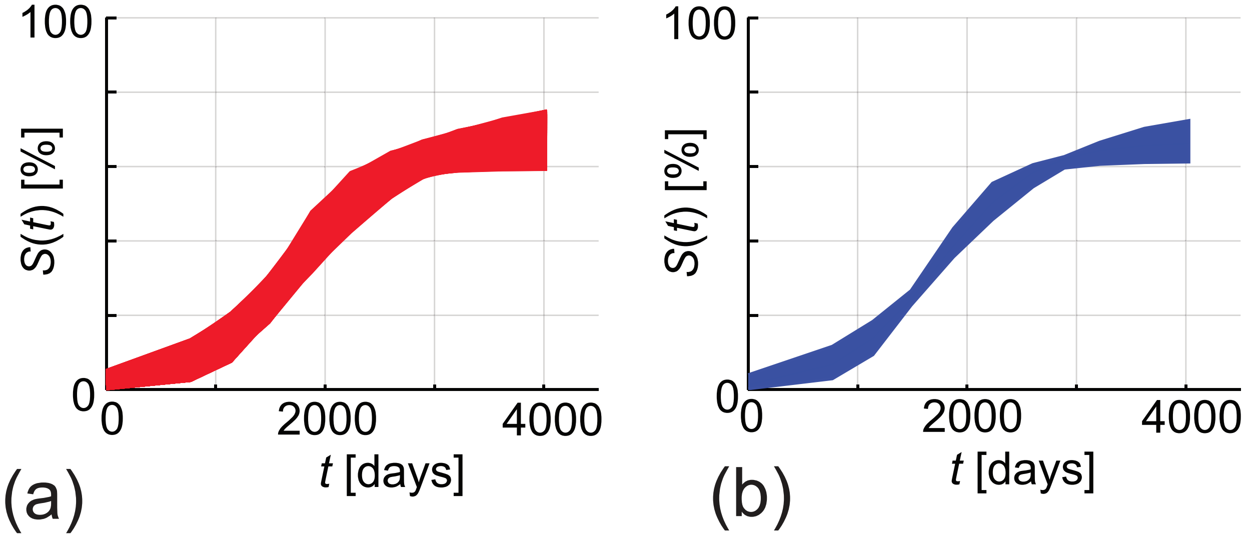


Figure S6: (a) prediction interval for the full likelihood (red curves). (b) approximate prediction interval constructed from profile likelihoods (blue curves).
